# Patterns and predictors of *Staphylococcus aureus* carriage during the first year of life; a longitudinal study

**DOI:** 10.1101/586032

**Authors:** Aylana Reiss-Mandel, Carmit Rubin, Ayala Maayan-Mezger, Ilya Novikov, Hanaa Jaber, Mordechay Dolitzky, Laurence Freedman, Galia Rahav, Gili Regev-Yochay

**Affiliations:** Infection Prevention & Control Unit, Sheba Medical Center, Ramat Gan, Israel; Sackler Faculty of Medicine, Tel Aviv University, Tel-Aviv, Israel; Gertner Institute, Tel Hashomer, Israel; Department of Neonatology, Sheba Medical Center, Ramat Gan, Israel; Maternity and Gynecology Center, Sheba Medical Center, Ramat Gan, Israel; Infectious Diseases Unit, Sheba Medical Center, Ramat Gan, Israel

## Abstract

**Objectives’:** To determine the patterns of *S. aureus* carriage in the first year of life, its determinants and dynamics of transmission between mothers and infants.

**Methods:** Prospective longitudinal cohort study of *S. aureus* carriage among mothers and their infants. Monthly screenings from pregnancy/birth through the first year of the infant’s life. Medical and lifestyle data was collected. Infant *S. aureus* carriage was detected by rectal and nasal swabs and maternal carriage by nasal swabs. Multivariate analysis and an NLMixed model were used to determine predictors of carriage and *S. aureus* persistence.

**Results:** 130 *S. aureus* carrier women and their 132 infants were included in the study. 93% of the infants acquired *S. aureus* sometime during the first year of life, 64% of them acquired the maternal strain, mostly (66%) during the first month of life. 70 women (52.50%) and 17 infants (14%) carried *S. aureus* persistently. Early acquisition of *S. aureus* carriage was associated with longer duration of initial carriage and was the most significant predictor of *S. aureus* persistence, while day-care center attendance was negatively associated with persistent carriage.

**Conclusions:** Early acquisition of *S. aureus*, mostly from the mother, is an important determinant of carriage persistence in infancy.

## Introduction

Asymptomatic carriage of *S. aureus* is common, with approximately 30% nasal carriage reported in the healthy population. Nasal *S. aureus* carriage has been shown to be an important source of transmission, as well as a significant source of endogenous infection (1, 2).

Risk factors of *S. aureus* carriage have been studied extensively in the adult population (3) and include age (3, 4), male gender (5–7), smoking (8), diabetes (9) and skin diseases, particularly atopic dermatitis (10). Longitudinal studies that reported carriage patterns, have found that 20% of adult healthy population are persistent carriers, typically of a single strain, 60% transient carriers and 20% are never carriers (11–13). Much less is known about early infancy carriage patterns or predictors of carriage.

Here, in this longitudinal study, we follow a cohort of infants born to *S. aureus* carrier mothers monthly, from birth until the age of one year, and observe the carriage patterns and carried strains, and their determinants during the first year of life.

## Methods

### Institutional Review Board (IRB) and patient consent

IRB approval was given by the local committee of the Sheba Medical Center. Written informed consent was given by the women for her and her newborn’s participation and a non-written approval of the other parent was also received.

### Study design, study period and study population

In this prospective longitudinal cohort study, pregnant women, at least 34 weeks of gestation, who visited the monitoring unit during screening hours, were recruited and screened for nasal and vaginal *S. aureus* carriage. Only women who were detected as nasal or vaginal *S. aureus* carriers, were enrolled and followed. Recruitment took place for 3 hours a week between February 2009 and March 2018 at the Sheba Medical Center obstetrics monitoring unit. The Sheba Medical Center is the largest tertiary center in Israel, with approximately eleven thousand births per year. Approximately 400 women visit the obstetrics monitoring unit monthly for reasons including overdue pregnancies (40+ weeks), breech fetal positioning, low levels of amniotic fluid, babies with outlying measurements, and monitoring of any pregnant woman who came to the emergency room for any reason. Within 48 hours of delivery, the mothers were rescreened with vaginal and nasal swabs. Concurrently, newborns were screened with nasal and rectal swabs. Data addressing demographic details (age, number of siblings, pet ownership and smoking status), medical history, including obstetric history and pregnancy complications, co-morbidity, medication and antibiotic use, previous hospitalizations and breastfeeding status as well as pregnancy and delivery details were collected via a questionnaire and from the electronic medical files. Screening was performed by the attending midwife, obstetrician or pediatrician at the delivery room or at the nursery.

Monthly follow-up visits from the age of 1m and until 12m of age were carried out by a study coordinator at the infants’ homes. During these visits mothers were screened with nasal swabs and children were screened with nasal and rectal swabs. Data addressing changing nutritional habits and medical events, including healthcare visits, antibiotic use, vaccination and hospitalizations were also collected.

### Laboratory methods

Nasal screening was performed using a cotton-tipped swab placed in Amies transport media (Copan innovation, Brescia, Italy). Swabs were streaked on CHROMagar *S. aureus* plates (HiLabs, Rehovot, Israel) within 24 hours and incubated for 24-48h at 35°C. Catalase and Staphylase (PASTOREX^®^ STAPH-PLUS, BioRad, Marnes-la-Coquette, France) were performed on suspected colonies to conclusively identify them as *S. aureus*. Cefoxitin agar disk diffusion test was used to detect methicillin resistant *S. aureus* (MRSA) according to the current clinical and laboratory standards institute (CLSI) protocol.

Genetic relatedness between mother and newborn strains were assessed by pulse field gel electrophoresis (PFGE) and spa typing. Maternal strain acquisition by the newborn was defined as acquisition of a *S. aureus* strain that was identical (by PFGE or spa typing) to his/her mother’s strain.

PFGE was done following the European HARMONY protocol (14). Briefly, digested DNA with *Sma*I was electrophoresed in 1% agarose gels for 21 hours with a ramped pulse time of 5 to 40 seconds using a CHEF DRII system (Bio-Rad Laboratories), using *S. aureus* NCTC 8325 as a reference. Genetic identity between strains was defined according to Tenover (15).

At least one strain from each pulsotype, and any strain where PFGE result was not available were Spa typed. Spa typing was performed by purifying the PCR product (Gene JET PCR DNA Purification kit, Fermentas) of the spa gene encoding protein A, using the primers 1517R: GCT TTT GCA ATG TCA TTT ACT G and 1095F: AGA CGA TCC TTC GGT GAG C. PCR products were Sanger sequenced by Hy Laboratories Ltd. (Rehovot, Israel), using BigDye terminator v1.1 Cycle Sequencing Kit (Applied Biosystems, Inc.) on the 3730xl DNA Analyzer with DNA Sequencing Analysis Software v. 5.4.Sequences were analyzed using the Fortinbras SpaTyper (http://spatyper.fortinbras.us/) and Ridom Spa Server (16). Genetic relatedness of strains was evaluated based on spa repeat patterns using the Tree and Network Inference module of Bionumerics Seven.

### Carriage patterns and definitions

Transient carrier mothers were defined as individuals who were colonized with *S. aureus* in less than 33% of available screenings. Persistent carrier mothers were defined as mothers who were colonized with *S. aureus* in at least 67% of the screenings available. Since most of the newborns acquired *S. aureus* within the first two months of life, using the above definitions would define many of the infants as persistent carriers, including those who only carried *S. aureus* for 2-3 months but were lost to follow up before the end of the year. We therefore used a more stringent idefinition for infant carriage: Persistent carriage of infants was defined as *S. aureus* carriage detected in at least 67% of the screenings available and also in at least 50% of the screenings from the second half year of life (age 6-12 months). Either rectal or nasal carriage, were considered as child *S. aureus* carriage.

### Statistical analysis

Descriptive data analysis was performed and Chi square test was used to examine the associations between categorical variables (i.e. mother’s and child’s carriage pattern). Spearman’s rho was calculated to evaluate the correlation between continuous variables (i.e. first month of *S. aureus* acquisition and duration of infection).

Initially, to explore which variable predicts persistent *S. aureus* carriage in the child, a univariate analysis was done on the following variables: sex, gestational age, birth weight, breastfeeding, pets, antibiotics in first year of life, skin infections, attendance at day care center (DCC), maternal carriage persistence, maternal carriage in first month of life, infant carriage in the first 2 months. Variables that were found to be associated with persistent infant carriage (p<0.2) in the univariate analysis were included in the multiple logistic regression model.

To determine the predictors for *S. aureus* carriage each month during the follow-up period, the altering demographic and clinical factors that independently predict *S. aureus* carriage in the following month were assessed. These factors included: infant’s carriage status in the preceding month, maternal carriage status in the preceding month, DCC attendance in the preceding month, antibiotic use in the preceding month, breastfeeding in the preceding month, and age. To account for the multiple measurements per subject in the longitudinal design, a non–linear mixed model (NLMIXED procedure) which fits a logit model was applied. Data were analyzed using SAS v9.4.

## Results

### Study population

Of all women approached, approximately 30% agreed to participate and take part in the monthly visits for the full year of follow-up. They were screened and signed an informed consent. Of the 671 women who were recruited, 136 were carriers of *S. aureus* in the nose or vagina at recruitment and were enrolled and followed in our study. Of these, 130 women and their 132 newborns completed at least 6 months of follow-up and were included in the final analyses. A total of 6043 swabs were collected from the mothers and children and 1887 *S. aureus* isolates were detected, 786 from the children and 1101 from the mothers.

Of the planned 12 monthly follow-up visits, 121 out of the 130 (93.1%) mother-child dyads completed at least 8 visits and 103 (79.2%) completed at least 10 visits. The 130 mothers included in the final analyses had a mean age of 34.2 years, (range 21 to 43, median 34) and a mean education level of 16.2 (+/−2.2) years. Thirty three women (25.3%) delivered their baby by Cesarian section (**Table 1**).

**Table 1.** Study Population Characteristics

| Variable |  | No. | % | Range | Mean |
| --- | --- | --- | --- | --- | --- |
| **Mother** |  | n=130 |  |  |  |
| Age (years) |  |  |  | 21 to 43 | 34.24 (+/- 4.71) |
| Education (years) |  |  |  | 12 to 20 | 16.20 (+/- 2.20) |
| Cesarean Section Birth |  | 33 | 26.83% |  |  |
| **Child** |  | n=132 |  |  |  |
| Gestational age |  |  |  | 34.57 - 42.14 | 39.93 (+/-1.48) |
| Birth weight (g) |  |  |  | 2030-4396 | 3310.41 (+/-496.12) |
| DCC attendance |  | 74 | 56.06% |  |  |
| (age at entry, months) |  |  |  | 3-12 | 7.1 (+/ 2.44) |
| Breastfeeding |  | 119 |  |  |  |
| (duration, months) |  |  |  |  |  |
|  | > 1 m | 114 | 86.36% |  |  |
|  | > 3 m | 104 | 78.79% |  |  |
|  | > 6 m | 75 | 56.82% |  |  |
| Antimicrobial use |  | 72 |  |  |  |
| months with use |  |  |  |  | 1.81 (+/ 1.25) |
| Physician visits | Ever during the year | 126 | 95.45% |  |  |
|  | Visited >2 times | 100 | 75.76% |  |  |
|  | Visited >3 times | 78 | 59.09% |  |  |
|  | Visited >5 times | 45 | 34.09% |  |  |
| Hospitalizations |  | 12 | 9.09% |  |  |

The children population was a normal birth cohort and children’s characteristics are described in detail in **Table 1**. Approximately half of the children (74, 46.1%) attended day care center (DCC) at some point during their first year of life, and of those, the median age of entry to DCC was 7 months (range 3-12 months). Most of the children were breastfed (n=119; 90.2%). Of these, 75 (63.0%) were breastfed for at least 6 months.

Health utilization during the first year of life was relatively high, with 72 (54.6%) children consuming at least one antibiotic regimen, 32 (24.2%) consumed at least two regimens and 15 children (11.4%) consumed more than 3 regimens during the follow up period. Nearly all of the children (n=125; 94.7%) had at least one episode of upper respiratory tract infection and 100 (75.8%) children visited their primary care physician more than twice during the year for non-routine vaccination visits. 12 (9.1%) children were hospitalized during the year (**Table 1**).

### Isolated Strains

Altogether, 119 clones were detected in 1887 bacterial isolates isolated from the 130 dyads over the course of the year. CC30 was the most frequently carried clonal complex in our sample, based on identification and grouping of spa typed strains. It was isolated 134 times; 9 different strains belonging to CC30 were isolated from the noses of 23 dyads. t3243 (CC22) was the most frequently isolated single strain, as it was carried by 10 dyads (**Table 2**). No clonal complex was found to be carried more commonly by persistent carriers than by transient or non-carriers.

**Table 2.**
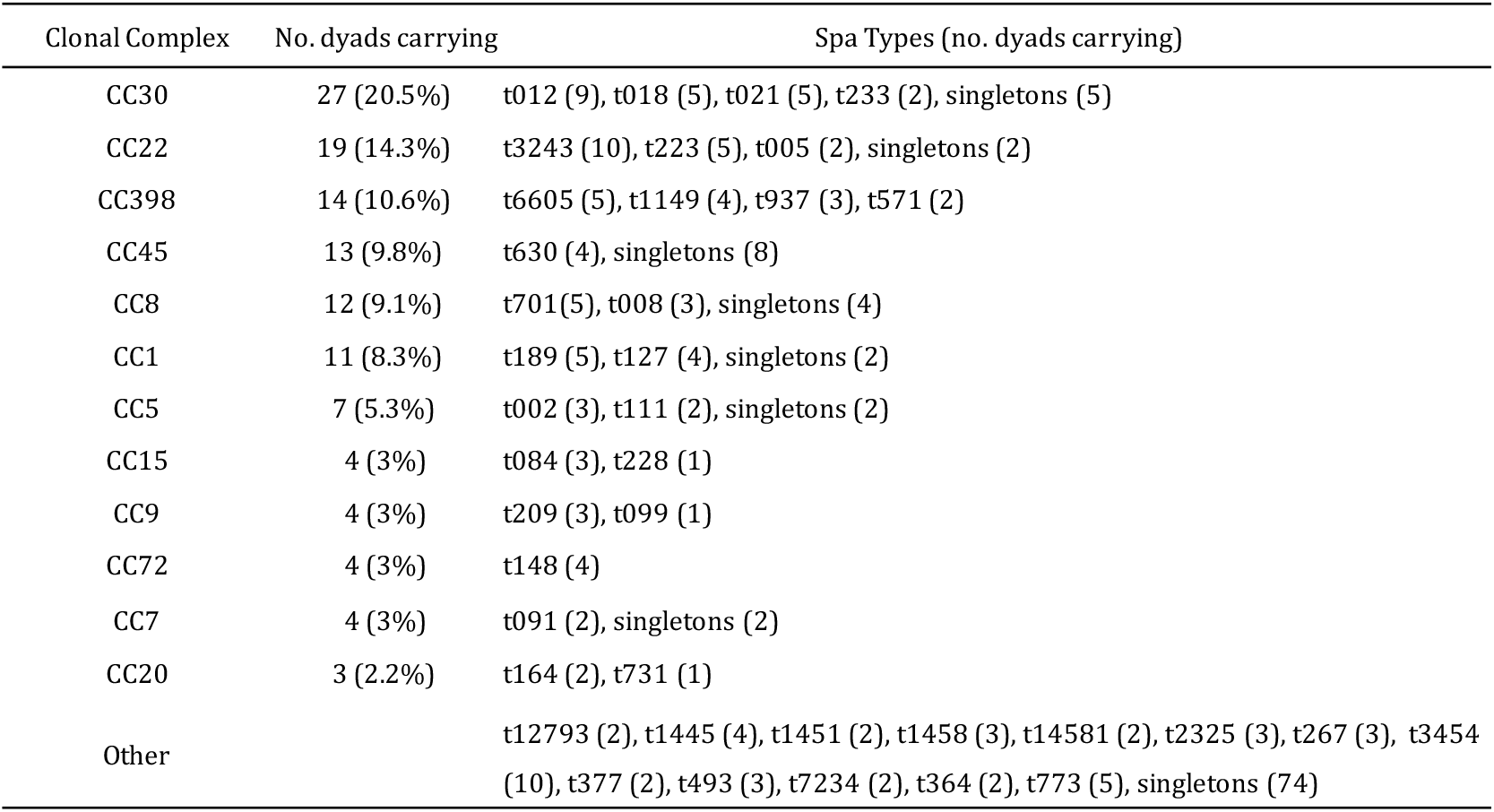
Most commonly carried strains

### Maternal S. aureus carriage patterns

At recruitment, 80 women carried *S. aureus* in the nose and 28 carried it in the vagina. 22 women carried *S. aureus* in both the nose and the vagina, and 17 of them carried the same strain in both sites. 57 women carried *S. aureus* in at least one site at both recruitment and immediately surrounding labor. In 54 cases (91.2%), the same strain was carried during gestation and labor, regardless of the carriage site.

Most of the participating women were defined as persistent carriers (n=70, 53.9%), while nearly a quarter were transient carriers (n= 32, 24.6%) and for an additional 28 women it was difficult to determine the pattern of carriage since they were carriers approximately 50% of the time (**Table 3**). Of the 70 persistent carrier women, 49 (70%) carried a single strain along all screenings during the year, as defined by PFGE or spa type, while 21 (30%) women carried a second strain at some point during the year. Nine of these women carried the secondary strain for only a month or two, after which the primary strain was again detected, while six women exchanged their initial strain with a second strain that was carried for most of the follow-up visits, and four women replaced their primary strain with a second strain that was carried for an extended period. (**Figure 1a**). No relationship was found between the length of carriage of a transient strain and its genetic relatedness to the persistent strain.

**Figure 1.**
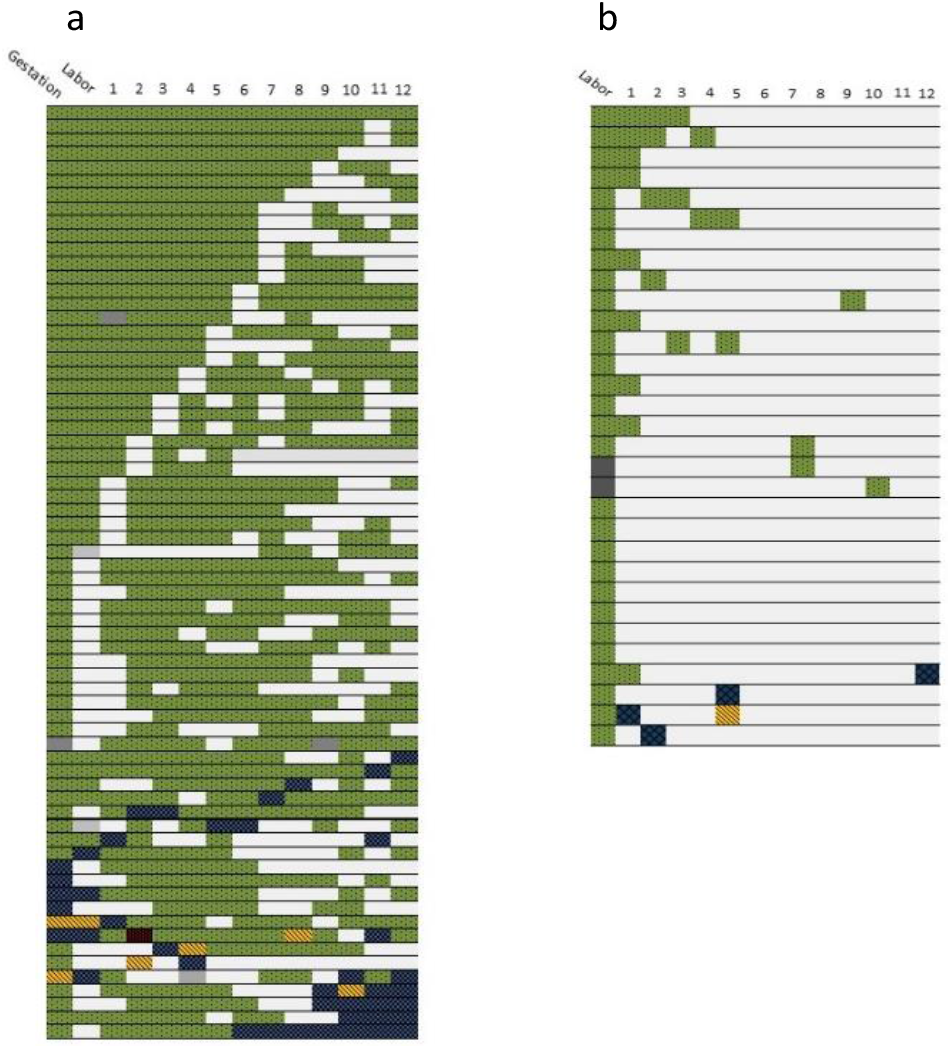
Patterns of carriage among mothers. (a) Carriage patterns of persistent carrier mothers (n=70), (b) carriage patterns of transient carrier mothers (n=32). Green strain indicates the main strain carried by the mother. Blue indicates the secondary strain carried by the mother. Yellow and red indicate third and fourth strains carried, when applicable. Grey indicates a missed screen or unidentified strain.

**Table 3.**
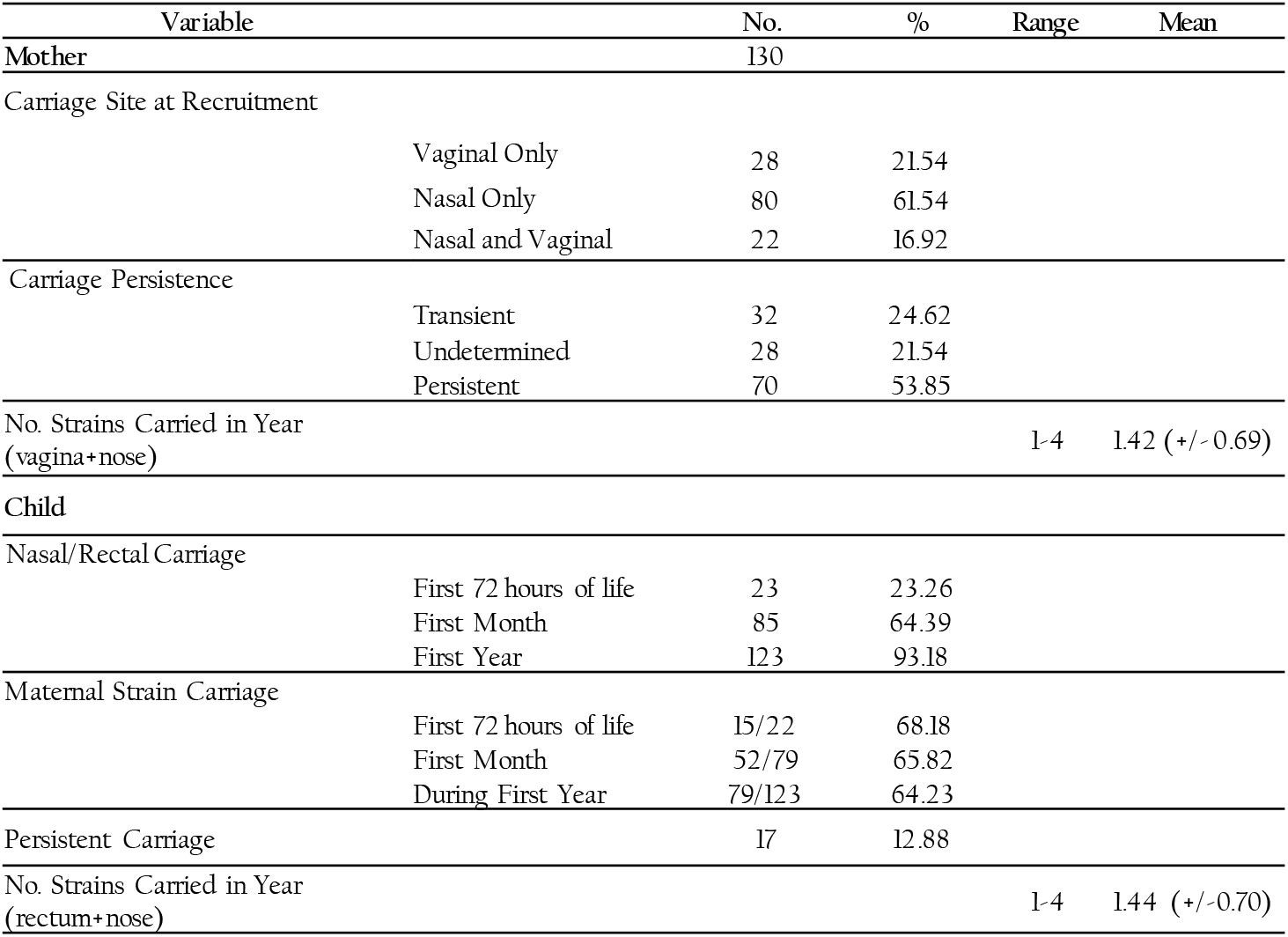
Maternal and infant carriage patterns and dynamics

Of the women defined as transient carriers (n=32), 15 (46.9%) never carried *S. aureus* during the year, apart from the initial screening when recruited during pregnancy. Of the transient carrier women who carried *S. aureus* in more than One visit, 5/15 (33.3%) carried two different strains in the different visits (**Figure 1b**).

### Infant S. aureus acquisition and patterns

Within the first year of life, 123 (93%) of the infants acquired *S. aureus*. Thirty (23% newborns acquired *S. aureus* in at least one body site (nose, rectum, ear or umbilicus) within the first days of life, before discharge from the hospital. Twenty three of them (76.7%) carried it in the nose or rectum. Initially, infant carriage was evenly distributed between nasal and rectal carriage. At 1 month, 84.2% were nasal carriers, and 47.4% were both nasal and rectal carriers. With time, rectal carriage prevalence decreased and the nose became the predominant site. (**Figure 2**). Over half of the infants (67) carried both a rectal and nasal strain in the same month at some point over the course of the year. The nasal and rectal strains were genetically identical in 85% (57/67) of the screens in which *S. aureus* was isolated from both sites.

**Figure 2.**
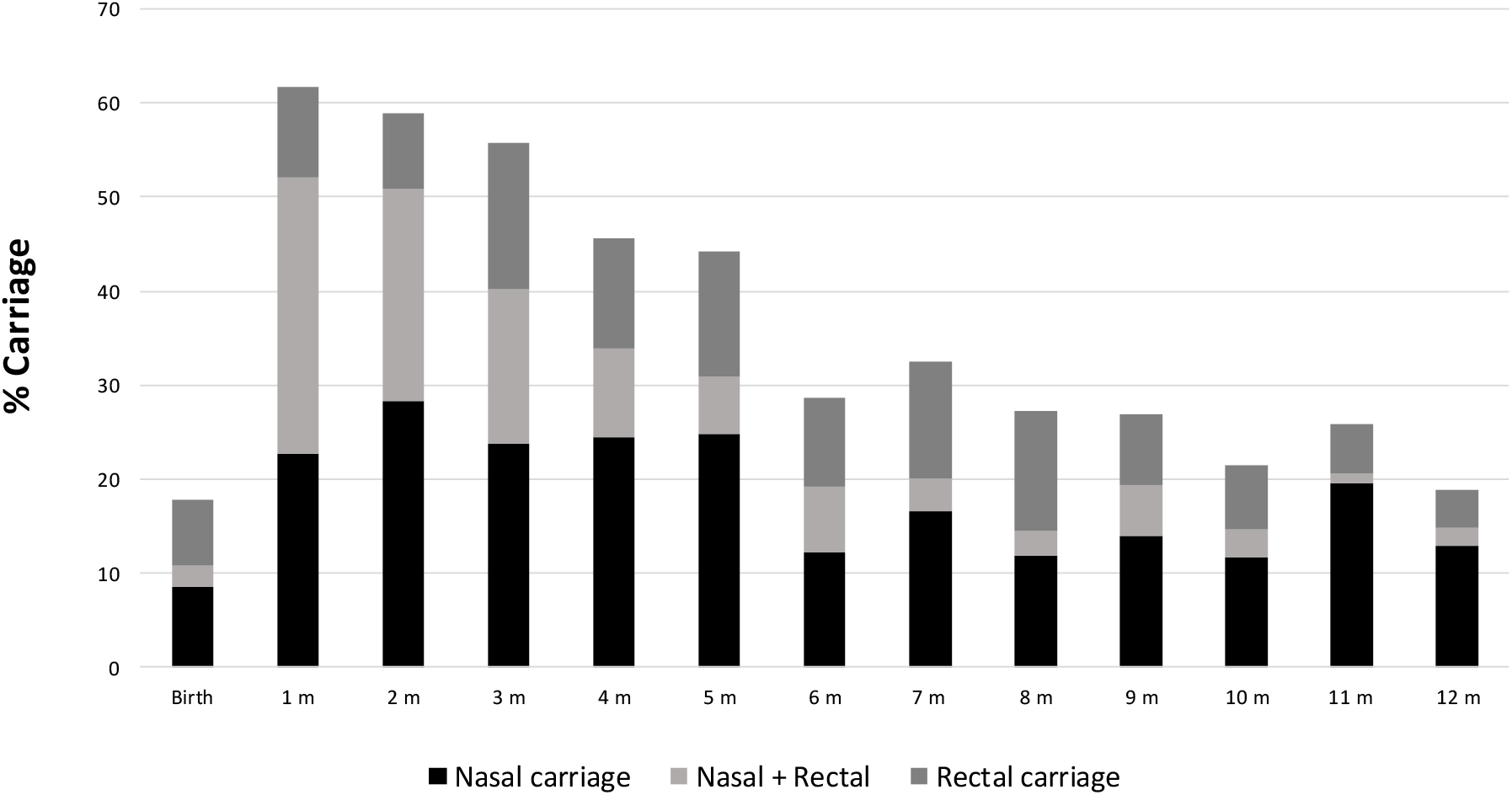
Infant nasal and rectal S. aureus carriage by month. Bars represent the percentage of all screened children who carried S. aureus nasally (black), rectally (dark grey) and both rectally and nasally (light grey) in a given month

**Figure 3.**
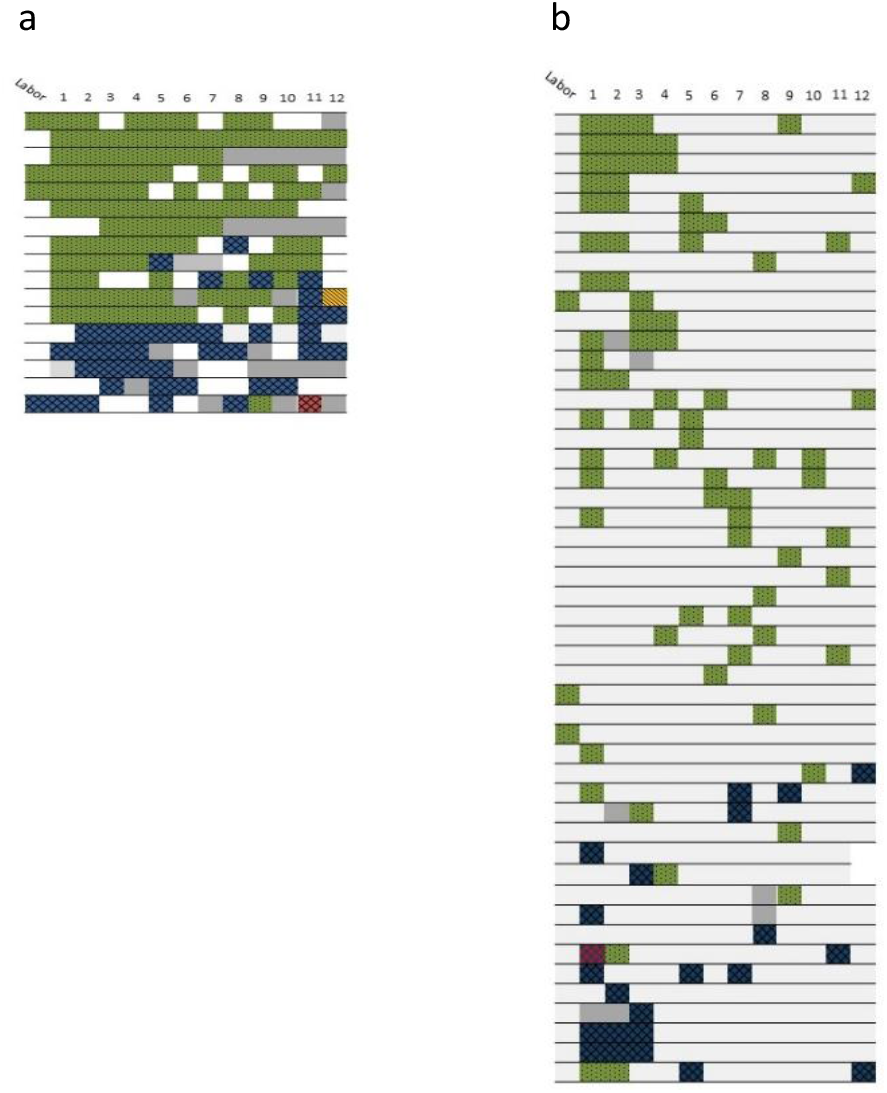
Infant carriage patterns (a) Carriage patterns of persistent carrier infants (n=17), (b) carriage patterns of transient carrier infants (n=52). Green strain indicates the main strain carried by the infant’s mother. Blue indicates the secondary strain, acquired by the child from a different source (c,d). Yellow and red indicate third and fourth strains carried, when applicable. Grey indicates a missed screen.

Of the 22 children that acquired *S. aureus* in the nose or rectum in the first days of life for whom strain data was available, 15 (68.2%) acquired the maternal strain (**Table 3**). On the 1^st^ month visit, 61.8% (n= 76) of infants were *S. aureus* carriers, 52 (72.2%) of them carried the maternal strain at this point.

None of the isolates carried by the mothers during the whole follow up period were methicillin resistant (MRSA). Two infants acquired non-maternal MRSA strains surrounding birth, during their stay in the hospital, but both replaced their strains in the next month, one with the maternal strain, and one with a different, unrelated strain.

Four patterns of carriage could be identified in the infants; (1) Persistent carriers (n=17, 13%), those who carried *S. aureus* at least 66% of screenings, but also at least 50% of the screenings in the 2^nd^ half of the year, (2) Never carriers, infants who did not acquire *S. aureus* during the whole study period (n= 9, 7%), despite the mother being a carrier on enrollment. (3) Transient carriers, infants who carried *S. aureus* for less than 34% of screenings (n= 55, 42%), and (4) a group of undetermined pattern (n=51, 38%) who did not meet the criteria for any of the above carriage patterns (**Table 3**).

Most persistent carrier infants carried a single strain over the course of the year, similar to the observation in the mother population. Over 70% (12/17) of these infants persistently carried the maternal strain (**Figure 4a**), while 5 (29.4%) carried a strain that was different than the strain isolated from their mother. In 4/5 of these cases, the mother never carried the infant’s persistent strain.

**Figure 4.**
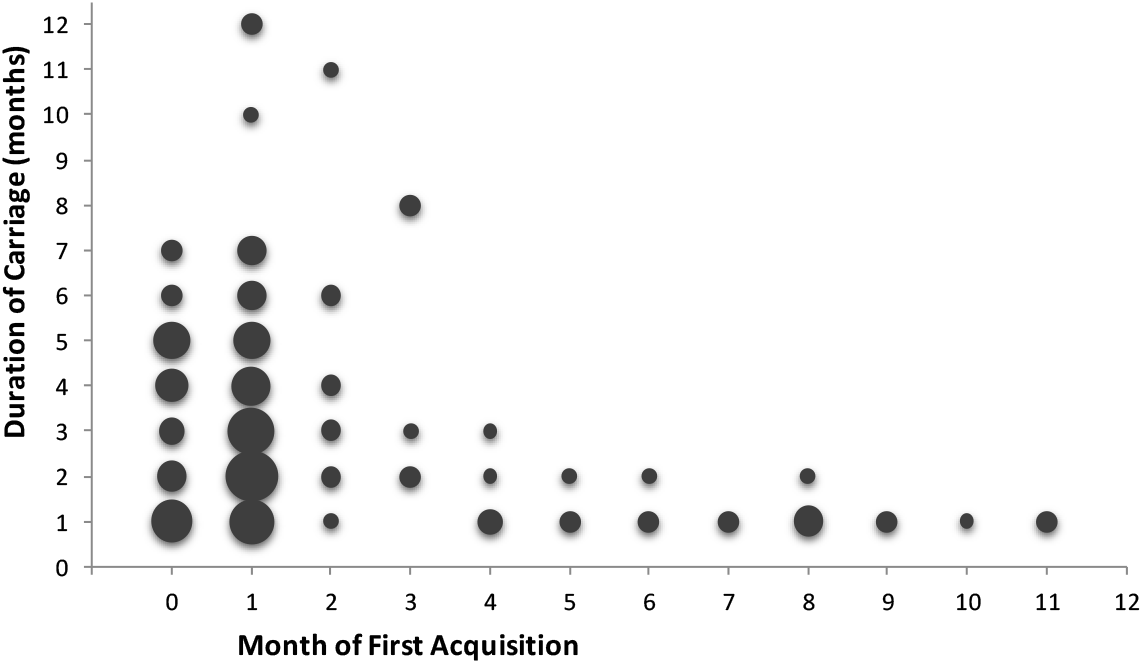
First acquisition of carriage vs duration of initial carriage. Circle size indicates amount of infants whose acquisition vs duration intersect at that point.

Of the nine infants who were never carriers, three had a persistent carrier mother. Three of the mothers of never carriers only carried *S. aureus* in the agina and not in the nose at recruitment and birth, and another five mothers carried *S. aureus* in the nose at recruitment but screened negative for nasal carriage at birth and/or the first month of follow up.

The most common pattern observed among mothers and infants was that in which the primary maternal strain was carried by both the mother and infant (n=76, 58%). In 28 cases (21.5%), the infant did not acquire the maternal strain at any point during the year.

No difference was observed between the number of strains carried by the mother and baby throughout the year. (**Table 3**). Most mother-infant dyads (85%) shared a strain at least once during the year.

Having a persistent mother was more common among persistent carrier infants (58.8%), compared to transient carrier infants (49.1%) or to never carrier infants (33.3%). Yet, having a persistent mother was not an independent predictor for infant carriage persistence, in a univariate analysis (p=0.38).

### Predictors of S. aureus carriage during the first year of life

The significant independent predictors for carriage in any given month were age, carriage of *S. aureus* in the previous month as well as maternal carriage in the previous month (**Table 4**).

**Table 4.**
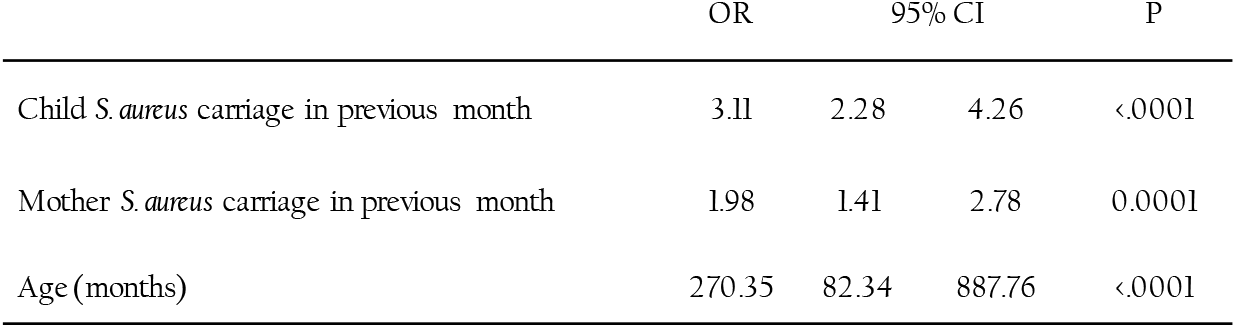
Independent predictors of child *S. aureus* carriage in each month

The major independent predictor for an infant to be a persistent carrier, was early acquisition of *S. aureus*, before the age of 2 months (**Table 5**). Furthermore, when assessing the association between the time of first *S. aureus* acquisition and the duration of carriage, we observed that the time of first acquisition predicts the duration of the carriage; The earlier the first acquisition, the longer it was carried (r=−0.3, p=0.0007) (**Figure 4**). While children who acquired the first strain during the first 2 months of life carried it on average for 3.68 (+/− 2.42) months (ranging from 1-12 months, median: 3), all of the children that acquired the first strain after the age of 4 months, had very short-lived carriage (range: 1-4 months, median:1).

DCC attendance was negatively associated with persistent *S. aureus* carriage in children (p=0.03). Sex, gestational age, maternal carriage persistence, birth weight, breastfeeding, pets, antibiotic use in the first year of life, breastfeeding, and skin infections were included in the univariate analysis but were not found to be significant and were not included in the multivariate analysis.

## Discussion

In this study, we followed maternal and infant *S. aureus* carriage throughout the first year of life. We found that most infants (93%) acquired S. aureus sometime during the first year, yet, most did not acquire *S. aureus* during birth, but acquired the maternal strain within the first month of life. Furthermore, we found that early acquisition of a *S. aureus* strain is the most significant predictor of long and persistent *S. aureus* carriage in the first year of life. These results point to the significant impact of maternal *S. aureus* carriage in the first months of a child life, on the infant’s carriage dynamics.

We show that over half of the mothers who were detected as carriers around labor, persistently carried *S. aureus*, mostly with single strain throughout the year. A quarter of the mothers were defined as transient carriers, of which most carried *S. aureus* in only one or two carriage events. In line with previous studies, we observed strain persistence in our healthy adult population, (12, 17). MRSA was not isolated from any mother at any point throughout the year. This is consistent with data from Israel, where community-acquired MRSA is not common (18).

In contrast to the extensive data on adult *S. aureus* carriage, not much has been reported on carriage patterns in early infancy. The duration of carriage, patterns and dynamics of carriage, or the predictors of carriage during infancy have been studied, but most studies did not continue past 6 months of age (19, 20) and those that followed the infants and mothers for an entire year only looked at 2-3 swabs during the course of the year (6, 21). Here, we screened the mothers and infants monthly, to obtain a full picture of their carriage patterns, as well as predictors of infant carriage in each month.

We previously reported that infants born to carrier mothers in the same cohort acquired their maternal strain in the first month of life (22). Here, we look at the full year and find that throughout the course of the year, 85% of infants acquired the maternal strain at least once, and that 12 of 17 persistent infants carried the maternal strain persistently. In line with observations by Jimenez-Truque et al, we observed that maternal carriage was a predictor of infant carriage at any given month, but, surprisingly, we did not find maternal persistence to be an independent predictor of infant persistence. Rather, it appeared that persistent carrier infants were likely to acquire their maternal strain early, within the first 1-2 months of life and that this early acquisition was the most significant predictor of persistent carriage, typically of a single strain, i.e. the maternal strain. Perhaps, in a larger sample size, maternal persistence would be a more significant predictor of persistent infant carriage. However, we did find that 33% of never-carrier infants were born to persistent mothers, this is low compared to persistent carrier infants, of whom 70% had a persistent carrier mother. Additionally, we and others have previously found a that infants carried or were infected by strains carried by both parents (18, 23, 24), and it is likely that looking at parental carriage patterns, as opposed to only maternal, would provide more evidence of an association between parent and infant persistence.

The sites of *S. aureus* carriage in early infancy have not been thoroughly studied previously. Here, we screened infants for both rectal and nasal carriage and observed that while prevalence of rectal and nasal carriage were almost equal surrounding birth, nasal carriage became the dominant site of carriage. We also observed that rectal and nasal strains were identical 85% of the time. In line with this, Lindberg et al (24) observed that strains isolated from rectal samples in the first 2 months of life were parental skin strains.

The role of the pathogen in determining the carriage pattern has been previously studied. No association between specific clones and carriage pattern was found by Muthukrishnan et al (13). Similarly, we did not find any correlation between specific strains (as determined by spa type) and carriage pattern in our population. Furthermore, we assessed whether among individuals that carried a secondary strain, the duration of carriage of the secondary strain would depend on the genetic relatedness to the primary strain, but did not find any statistically significant relation.

Previous findings by our group show that *S. aureus* carriers display a tolerogenic immune response to their own strain (25) and it is known that host-bacterial interactions in early life help shape the developing immune system and the commensal microbiome for years to come. These results, where we show that early acquisition of a strain predicts longer carriage during the first year of life, are consistent with the idea of a tolerogenic response to early acquisition of *S. aureus*, though long-term follow-up into late childhood or even adulthood are required to determine the implication of early *S. aureus* acquisition.

Our study has several limitations. Although this study was large and comprehensive, a larger population, possibly including follow up with non-carriers, or longer follow up time could provide greater statistical power to some predictors and correlations. Screening of other family members could also provide more insight and a more comprehensive picture of the dynamics of *S. aureus* carriage within the family and not only between the mother and baby.

Additionally, our results on genetic identity depend solely on the evolution of the spa gene.

As *S. aureus* carriage plays such a significant role in the dynamics of infection, understanding the initial acquisition is vital. This study is the first to show such a prominent role of early strain acquisition in both the carriage duration and pattern, as well as the intimate dynamics of *S. aureus* carriage between mothers and babies.

## Acknowledgments

We would like to thank Dr. Eyal Leshem who was involved in the early stages of the project. Yael Beker-Ilany who was devoted to the women and the children and carried out the monthly visits along most of the study years. We would like to thank the helpful Sheba Medical Center monitoring unit team and the nursery team.

This study was funded by the Chief Scientist, Ministry of Health, Israel (Grant <u>3-00000-5622</u>) and the Israel Science Foundation (1590/09, 1658/15).

---

Table 5. Independent predictors of persistent *S. aureus* carriage in infants

